# SERBP1 licenses activation of the integrated stress response at collided ribosomes

**DOI:** 10.64898/2026.11.26.769417

**Authors:** Zhen Ye, Shuhan Yang, Timo Denk, Toru Suzuki, Takuya Tomita, Carla M Engel, Matthias Thoms, Sihan Li, Satoshi Hashimoto, Chisato Kikuguchi, Alexander Landis, Yumiko Isobe, Kiyoshi Yamaguchi, Yoichi Furukawa, Chloe S Palma-Chaundler, Christian Blum, Stefan Krebs, Jia-Xuan Chen, Yasushi Saeki, Roland Beckmann, Julian Stingele, Toshifumi Inada

**Author notes:** These authors contributed equally (Zhen Ye, Shuhan Yang, Timo Denk, Toru Suzuki).

## Abstract

Activation of the integrated stress response by GCN2 lets cells adapt to translational stress. GCN1 recruits GCN2 to collided ribosomes, but the ribosome-associated factors that couple collisions to GCN2 activation remain incompletely defined. Here we identify SERBP1, previously characterized as a factor that excludes surveillance machinery from ribosomes, as a component of the GCN2-dependent response. Correlation analysis of genetic screens linked SERBP1 to GCN1 and GCN2, and loss of SERBP1 impaired GCN2 and eIF2 alpha phosphorylation, stress-response gene induction and translational repression after messenger RNA damage and elongation stress. Biochemical and structural analyses revealed a multivalent ribosome-bound platform formed with RACK1 and GCN1. Conversely, loss of SERBP1 or GCN1 increased ZNF598-dependent ribosomal ubiquitination and quality control independently of GCN2 kinase activity.

## Introduction

Translation is continuously challenged by damaged mRNAs, limited substrate availability and intrinsic barriers to elongation (*1*). As a consequence, elongating ribosomes can stall, and if stalling persists, be encountered by trailing ribosomes, generating collided assemblies recognized by dedicated surveillance pathways. Structural studies revealed a characteristic 40S-40S interface that provides a preferred substrate for the ubiquitin ligases Hel2 in yeast and ZNF598 in mammals (*2, 3*). Hel2- or ZNF598-dependent ubiquitination of small-subunit proteins promotes ribosome resolution and downstream mRNA and nascent-chain quality control through the ribosome-associated quality control-trigger (RQT) complex in yeast and the related ASCC machinery in mammals (*4–9*). Thus, ribosome collision provides a conserved signal for recognizing aberrant translation complexes.

Collided ribosomes are not committed exclusively to destructive quality control. They also activate the GCN2-dependent integrated stress response (ISR) and the ZAKα-dependent ribotoxic stress response (RSR), establishing ribosome collisions as platforms for cellular stress signalling (*10*). GCN1 provides a central connection between ribosomal state and GCN2 activation. Structural analysis showed that yeast Gcn1 spans stalled and collided ribosomes within a disome (*11*). More recently, reconstitution of the response to amino acid deprivation demonstrated that GCN1 functions as a disome sensor that recruits GCN2 to collided ribosomes, where ribosomal interactions promote GCN2 activation and stable association with the collision (*12*). Cellular RNA damaged by UV or reactive metabolites can engage the same pathway (*1, 10, 13*). Similarly, incorporation of the chemotherapeutic nucleoside analogue azacytidine (azaC) into mRNA stalls translating ribosomes and activates GCN1-GCN2 signalling, whereas RNF25-dependent ubiquitination of eS31 restrains excessive GCN2 activation (*14*).

The 40S-associated WD-repeat protein Asc1/RACK1 occupies a central position at the intersection of ribosomal surveillance pathways. Asc1 promotes translational arrest and no-go decay in yeast (*15*) and lies at the collision interface recognized by Hel2 (*3*), whereas mammalian RACK1 contributes to ZNF598-dependent quality control (*6*) and ZAKα signalling (*16*). Whether this shared ribosomal surface also supports GCN1-dependent recruitment and activation of GCN2, and how such a role might be coordinated with competing quality-control pathways, remains unclear.

SERBP1 is a conserved RNA- and ribosome-associated protein with an established role in ribosome dormancy. SERBP1 occupies the mRNA channel of non-translating mammalian 80S ribosomes together with eEF2 (*17, 18*) and promotes formation of dormant 80S ribosomes following mTORC1 inhibition (*19*). Additional cryo-EM structures revealed SERBP1-bound dormant states containing eEF2, eIF5A or tRNA (*20*). More recently, structural and biochemical analyses have extended the role of SERBP1 beyond dormant ribosomes, showing that SERBP1 also acts at translationally active ribosomes. Here, the C-terminal FPxL motif of SERBP1 binds RACK1 at a site overlapping the RACK1-binding motif of ZAKα, thereby antagonizing collision-dependent ZAKα activation (*16*). Proteomic analysis further showed that SERBP1 is abundant on translating ribosomes, where it broadly limits ribosome association of factors involved in ribosome-associated quality control (RQC), RNA decay and selective 40S degradation (*21*). SERBP1 loss increased association of ZNF598 and ASCC components with polysomes and promoted RNF10-RIOK3-dependent 40S degradation. These findings establish SERBP1 as a negative regulator of multiple ribosome interactions, but whether it also promotes recruitment of specific factors to collided ribosomes remains unknown.

Here, we identify a positive signalling function of SERBP1 at collided ribosomes. Unbiased genetic analysis connects SERBP1 with GCN1 and GCN2 in a functional network, and structural and functional analyses reveal a multivalent SERBP1 platform that engages GCN1 and supports GCN2 activation. Conversely, loss of SERBP1 or GCN1 enhances ZNF598-dependent ribosomal ubiquitination and RQC independently of downstream GCN2 activity. We propose that the SERBP1-GCN1 platform converts a physical collision into a GCN2-signalling-competent state while restricting engagement of RQC.

## Results

### SERBP1 promotes GCN2 activation

The GCN2-dependent ISR drives cytotoxicity following azaC-induced mRNA damage, and loss of GCN1 or GCN2 consequently confers strong resistance to azaC (*14*). To identify genes with a similar phenotype, and thus potentially unrecognized components of this pathway, we sequenced additional time points across our previous azaC CRISPR screens (*14*) (fig. S1a and table S2), enabling us to construct a gene network based on pairwise Spearman correlations (Fig. 1a, data S1 and table S3). SERBP1 showed a strong positive correlation with GCN1 and EIF2AK4, which encodes GCN2 (Fig. 1a). Similar to loss of GCN1 or GCN2, loss of SERBP1 consistently conferred azaC resistance across most acute and chronic azaC screens in HAP1 WT, RNF25- and ZNF598-knockout (KO) cells (Fig. 1b).

We confirmed this phenotype in clonal SERBP1-KO cells, which were more resistant to azaC than control cells, to a similar extent as GCN2-deficient cells (Fig. 1c, fig. S1b). Combined loss of SERBP1 and GCN2 did not further increase resistance relative to the individual KO cells, consistent with both proteins acting in the same pathway (Fig. 1c, fig. S1b). Moreover, SERBP1 loss suppressed the enhanced azaC sensitivity caused by GCN2 hyperactivation in RNF25-KO cells (*14*) (Fig. 1d, fig. S1b). Together, these data implicated SERBP1 in the GCN2-dependent ISR.

Consistent with this conclusion, phosphoproteomic analysis followed by pathway enrichment analysis indicated reduced GCN2 kinase activity in SERBP1-deficient cells following azaC treatment (fig. S1c, table S4). Immunoblotting showed reduced phosphorylation of GCN2 and eIF2α in SERBP1-KO cells, both in the absence and presence of azaC treatment (Fig. 1e). Accordingly, SERBP1 loss attenuated azaC-induced upregulation of ISR target genes, as measured by RNA-seq (Fig. 1f, table S5), and repression of global protein synthesis, as measured by O-propargyl-puromycin (OPP) incorporation (Fig. 1g). Both effects were comparable to, although slightly less pronounced than, those of GCN2 loss. Together, these data establish that SERBP1 is critical for GCN2-dependent ISR activation following azaC-induced mRNA damage.

**Figure 1.**
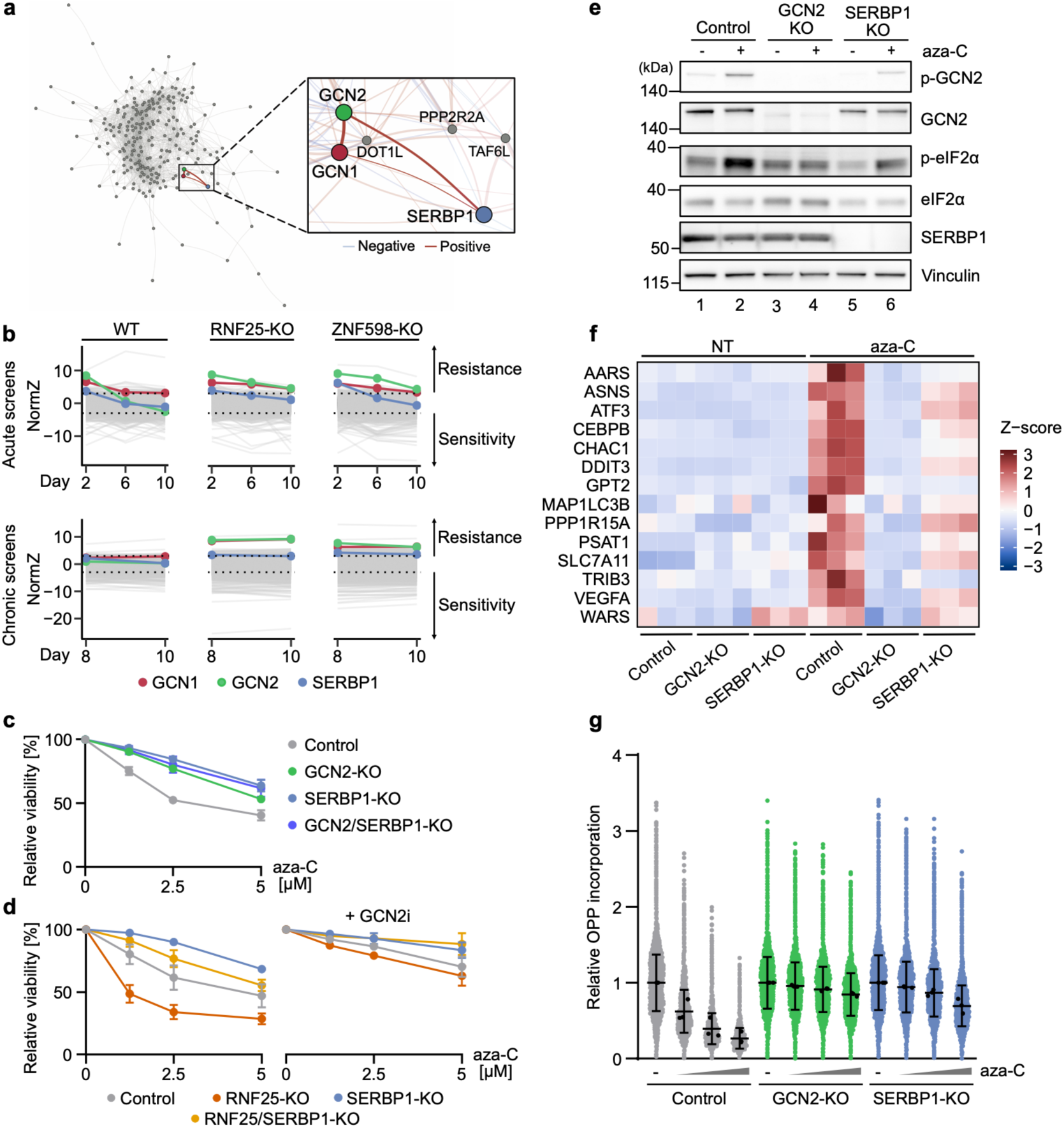
SERBP1 promotes GCN2-dependent ISR activation following azaC treatment. **(a)** Gene-network analysis of genome-wide CRISPR/Cas9 screens performed in HAP1 WT, RNF25-KO, and ZNF598-KO cells for genetic determinants of azaC sensitivity. Genes are positioned according to correlations between their phenotypic profiles across the analyzed screens. Enlarged view highlights the region of the network containing SERBP1, GCN1 and GCN2. Red and blue edges in the rectangle indicate positive and negative correlations, respectively, with edge width proportional to |ρ|. **(b)** Normalized Z-score trajectories across acute and chronic CRISPR/Cas9 screens. SERBP1, GCN1 and GCN2 are highlighted, while gray lines represent the trajectories of all other genes at the indicated time points. Dotted lines indicate thresholds at -3 and +3, corresponding to resistance and sensitivity hits. **(c-d)** Cell viability of HAP1 control and KO cells treated with the indicated concentrations of azaC in the presence or absence of GCN2 inhibitor (GCN2i; A-92, 1 µM) for 48 h. Values represent the mean ± SD of 3 independent biological replicates, normalized to the untreated condition for each cell line within each replicate. **(e)** Representative western blot analysis of HAP1 control, GCN2-KO and SERBP1-KO cells treated with or without 20 μM azaC. **(f)** Heatmap showing Z-scored expression values of ATF4 target genes (*22*) for HAP1 control, GCN2-KO and SERBP1-KO cells treated with or without 20μM azaC. **(g)** OPP incorporation assay measuring global translation rates in HAP1 control and KO cells treated with increasing concentrations of azaC.

### SERBP1 promotes the recruitment of GCN2 to collided ribosomes

To dissect how SERBP1 supports GCN2 activation at collided ribosomes, we sought an experimental model that avoids the pleiotropic effects of azaC, including DNA damage. We therefore first tested whether SERBP1 is also required for GCN2 activation following acute elongation stress. Indeed, anisomycin (ANS), like azaC, induced robust GCN2 and eIF2α phosphorylation in control cells, whereas both responses were strongly attenuated in SERBP1-KO cells, similar to GCN1 deficiency (Fig. 2a). Following ANS treatment, SERBP1 loss selectively impaired GCN2 and eIF2α phosphorylation while JNK and p38 activation remained robust (Fig. 2b). Consistent with the effects observed following azaC treatment, transcriptome analysis showed attenuated induction of ISR-associated genes, including ATF4, CHAC1, ASNS and SESN2, in SERBP1-deficient cells following ANS treatment (Fig. 2c). Quantitative proteomic analysis showed no widespread alteration in the abundance of small- or large-subunit ribosomal proteins in SERBP1-deficient cells under basal or ANS-treated conditions (fig. S2b). Thus, the ISR activation defect upon loss of SERBP1 is unlikely to result from a gross change in ribosome composition.

We next examined whether SERBP1 loss affects the composition and interactome of collision-associated ribosomes. For this, cytoplasmic extracts from DMSO- or ANS-treated cells were resolved by sucrose-density-gradient centrifugation, and disome fractions were analysed by immunoblotting and quantitative DIA mass spectrometry (Fig. 2d). At the low dose used, ANS did not substantially alter overall disome abundance, but GCN1 redistribution into the disome fraction indicated collision formation (Fig. 2d, fig. S2a,c). Quantitative mass spectrometry showed reduced association of eIF5A with disomes after ANS treatment, while EDF1, ZNF598, GCN1 and GCN2 were enriched in WT cells (fig. S2c; data S2). In disome DIA-MS analysis, GCN2 abundance increased 2.03-fold after ANS treatment in WT cells, compared with 1.006-fold in SERBP1-KO cells, relative to the respective DMSO-treated controls (fig. S2c; data S2). Notably, loss of SERBP1 abrogated ANS-induced recruitment of GCN2 to disomes (Fig. 2e; fig. S2c). In contrast, recruitment of GCN1 and other collision sensors, including ZNF598 and EDF1, was retained in SERBP1-deficient cells. Thus, GCN1 association with disomes is not sufficient for efficient GCN2 recruitment under these conditions: SERBP1 enables collided ribosomes to support GCN2-dependent signalling.

**Fig. 2.**
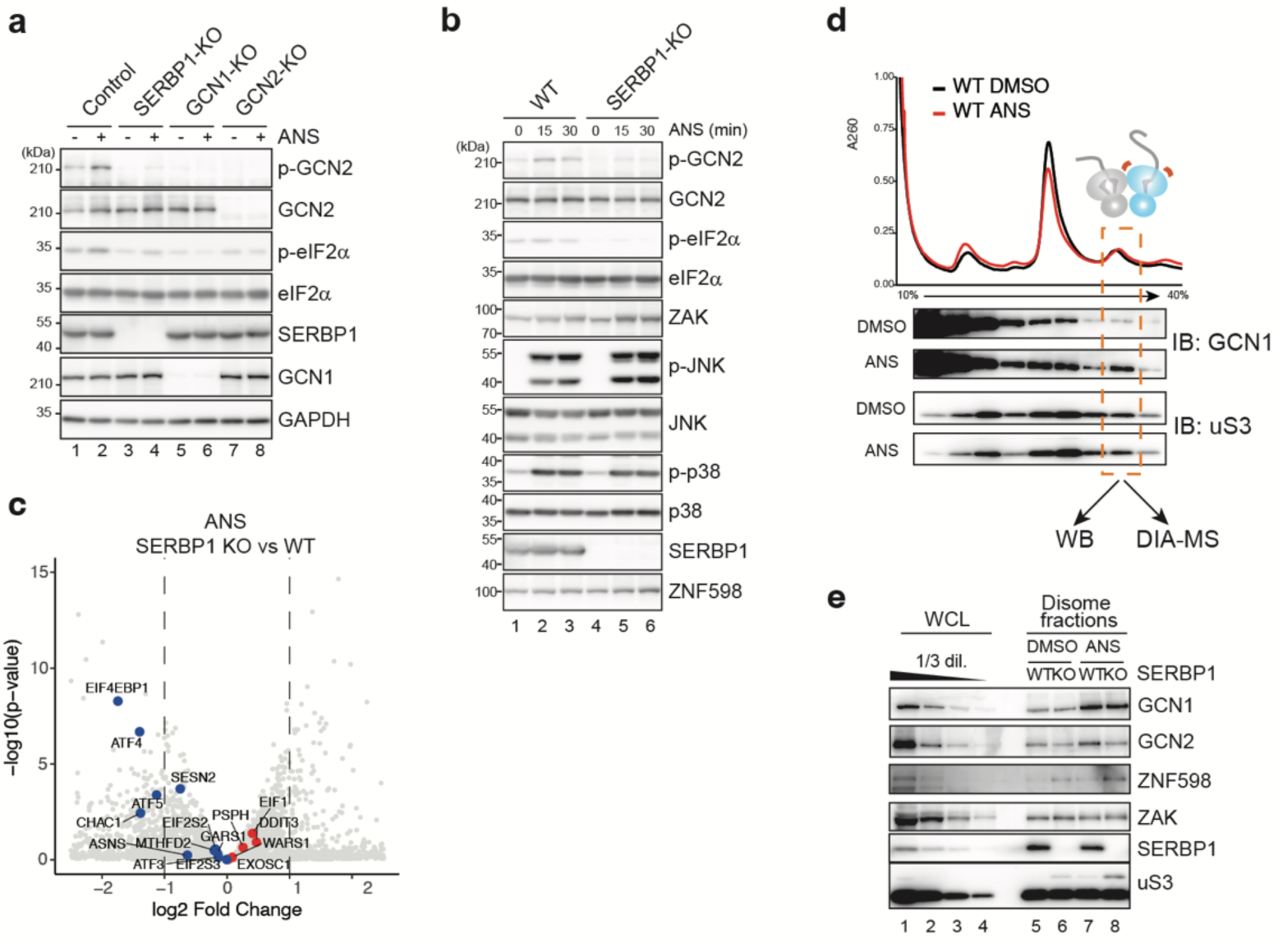
SERBP1 promotes assembly of a GCN1-GCN2 signalling platform on collided ribosomes. **(a)** Immunoblot analysis of GCN2-ISR activation in control, SERBP1-KO, GCN1-KO and GCN2-KO cells following treatment with anisomycin (ANS). Phosphorylated and total GCN2 and eIF2α were analysed together with SERBP1 and GCN1. GAPDH serves as a loading control. **(b)** Time-course analysis of ISR and RSR signalling in WT and SERBP1-KO HEK293T cells following ANS treatment for the indicated times. Phosphorylation of GCN2, eIF2α, JNK and p38 was analysed by immunoblotting, together with total proteins and the indicated controls. **(c)** Volcano plot showing differential gene expression between SERBP1-KO and WT cells following 15 min ANS treatment. The x axis indicates log2 fold change and the y axis indicates −log10(P value). Selected ISR-associated genes are indicated. **(d)** Isolation and analysis of collision-associated ribosome fractions. Cytoplasmic extracts from DMSO- or ANS-treated cells were separated by sucrose-density-gradient centrifugation. Absorbance profiles at 260 nm are shown above immunoblot analysis of gradient fractions for GCN1 and uS3. Fractions corresponding to the disome region (orange dashed box) were pooled for immunoblotting and DIA mass-spectrometric analysis. **(e)** Immunoblot analysis of whole-cell lysates (WCL) and purified disome fractions from WT and SERBP1-KO cells treated with DMSO or ANS. GCN1, GCN2, ZNF598 and ZAKα were analysed together with SERBP1. uS3 serves as a ribosomal marker. Serial dilution of WCL is shown for comparison.

### SERBP1 forms a multivalent ribosome-associated platform with GCN1 for GCN2 activation

To understand how SERBP1 cooperates with GCN1 to promote GCN2 activation, we first sought to further characterize its engagement with actively translating ribosomes on a molecular level. Therefore, we prepared an in extracto lysate sample (*23, 24*) from high dose ANS treated FreeStyle 293-F cells and subjected it to cryo-EM single particle analysis. After 3D classification of ribosomal particles, the most abundant state (∼42%, fig. S3) represented decoding 80S containing P/P A/T tRNAs and eEF1A (2.47 Å overall, Fig. 3a-c, fig. S3c and S4, table S1). On these particles we observed ANS at its characteristic position in the peptidyl transferase center (PTC) (*25, 26*) as well as SERBP1 forming multiple contacts with the ribosome at both the small and large subunit (Fig. 3a-d, fig. S5a). SERBP1 residues 288–408 span the 40S head, establishing three major ribosomal interaction sites (Fig. 3e,f). At the 40S beak, residues 288– 328 of SERBP1 extend from eS10 across eS12 towards eS31, where they form a β-augmentation interaction that anchors SERBP1 to the ribosome (Fig. 3e(I),f and figs. S5b,c and S6a,b). Thus, SERBP1 is positioned in proximity to RNF25’s key target residue eS31-K113, whose ubiquitination restrains GCN2 activation (*14, 27*). SERBP1 (A348-G360) further protrudes towards uS10 binding to a hydrophobic pocket and implementing another beta-sheet augmentation (Fig. 3e II, fig. S5d-f, S6a and b). This differs from ribosome preparations lacking SERBP1 where instead an extended alpha helix was observed for this region of uS10 (*28*) (fig. S5g). Lastly, the uS10 binding site is connected to the very C-terminus of SERBP1 via a positive stretch (363-387aa) that we did not observe and is likely flexible but would be close to the mRNA entry potentially interacting with the mRNA prior to entering the ribosome. The SERBP1 C-terminus itself (A393-A408) is binding RACK1 via its FPxL motif (404-FPAL-407) as observed before (*16*) (Fig. 3e III, f, fig. S6a and b). The very N-terminus of SERBP1 (G5-P27) on the other hand is positioned on the 60S, binding to eL27, eL30 and eL34 as previously postulated (*29*) (Fig. 3g, fig. S6a and b). Overall, our cryo-EM data show that SERBP1 forms a multivalent interaction network on the ribosome that physically links both ribosomal subunits and covers multiple key sites relevant in different surveillance pathways on the 40S (Fig. 3f and g, fig. S6a and b).

Having established the interaction of SERBP1 with active ribosomes we next investigated potential physical links to GCN1 and GCN2. The positioning of GCN1 on ribosome collisions based on in-situ cryo-EM data (*30*) infers close proximity of GCN1 to SERBP1 as the former extends over the small subunits of both stalled and collided 80S (Fig. 3h). Therefore, we performed AlphaFold 3 (*31*) (AF3) multimer predictions using GCN1 but also GCN2 whose ribosome-bound state remains structurally uncharacterized (fig. S6c and d). Two distinct GCN1– SERBP1 interaction sites were predicted with high confidence (Fig. 3f, 3h I and II). At the first interface, the N-terminus of SERBP1 was predicted to engage an internal part of GCN1 (∼1790– 1910 aa), positioned near the central protuberance on the stalled 80S (Fig. 3f, 3h I). At the second interface, a part of SERBP1 containing the uS10 interaction site (Fig. 3e II and f) was predicted to bind the N-terminus of GCN1 (∼1–127 aa). For GCN2, the C-terminus of SERBP1 (∼390–408 aa), containing the FPxL motif, was predicted to insert into the histidyl-tRNA synthetase-like (HRS) domain of GCN2 (Fig. 3f and i, fig. S6d).

To validate the predicted interfaces, we generated structure-guided SERBP1 mutants and tested their ability to complement the loss of endogenous SERBP1 in cells. Re-expression of WT SERBP1 restored ANS-induced GCN2 and eIF2α phosphorylation in SERBP1-KO cells, as expected (Fig. 3j-l). In contrast, mutation of the C-terminal FPxL motif (FPXL-3A, FPXL-5A) or the predicted GCN1-(GCN1-N, GCN1-M, GCN1-N&M), GCN2-(GCN2-HRS) and ribosome-interaction surfaces (DK346-G360) impaired GCN2 activation (Fig. 3j-l). Furthermore, GCN1- and ribosome-interaction surfaces mutants did not alter the association of GCN1 nor GCN2 with ribosomes (Fig. 3m).

Together, these data support a model in which ribosome-bound SERBP1 utilizes some of its ribosome binding stretches to multivalently assemble a joint platform with GCN1 that recruits and activates GCN2. Two copies of SERBP1, one on the stalled and another one on the colliding ribosome, could cooperate in this process.

**Figure 3.**
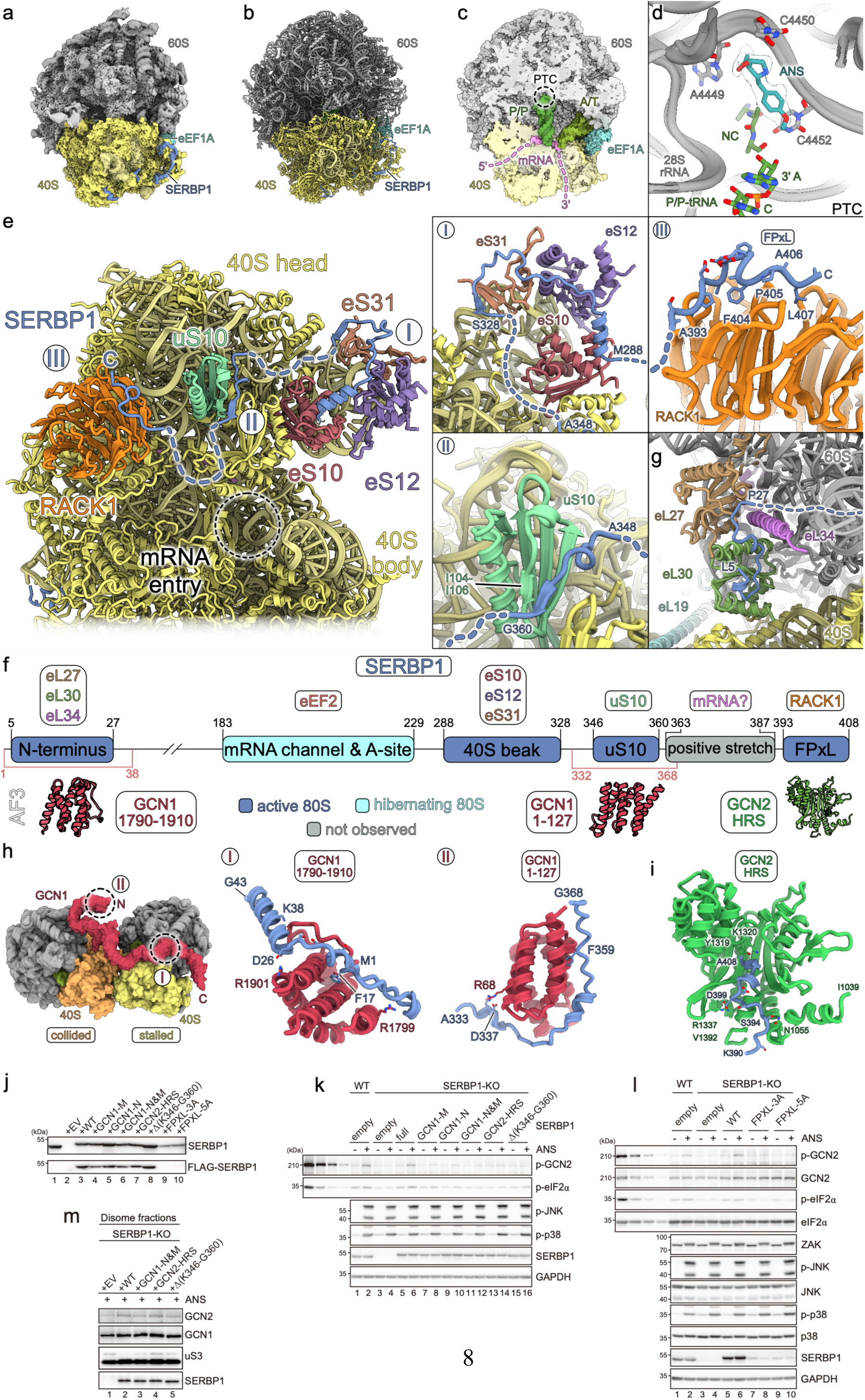
SERBP1 forms a multivalent ribosome-associated platform for GCN1-GCN2 activation. **(a-b)** Local resolution filtered composite cryo-EM density (a) and molecular model (b) of the SERBP1 and ANS bound human 80S. **(c)** Cut-through view of the molecular model in (b) shown as surface representation highlighting the translational state. **(d)** Close up of the PTC with density for ANS shown. **(e)** SERBP1 contact sites with the 40S head. Overview (left) as well as close ups of the individual contact sites at the 40S beak (I), ribosomal protein uS10 (II) and RACK1 (III). **(f)** Simplified schematic overview of SERBP1 with its distinct interaction motifs annotated. On top interactions with ribosomal proteins and translation factors are shown. On the bottom potential interaction sites with GCN1 and GCN2 based on AF3 predictions (see fig. S6c and d) are indicated. **(g)** Close up of the SERBP1 N-terminus binding site at the 60S subunit. **(h)** Putative visualization of GCN1 binding to a collided ribosome. The model is based on GCN1 bound to collided ribosomes from in-situ cryo-electron tomography density (EMD-19242) (*30*) and the shown surface representation has been generated by using PDB:9RPV (*16*) and the AF2 database GCN1 prediction. Predicted SERBP1 binding sites are indicated. Close-ups of the two predicted binding sites of SERBP1 to GCN1 are shown (I and II) (see fig. S6c). **(i)** Close up of the predicted interaction of the SERBP1 C-terminus with the GCN2 HRS domain (see fig. S6d). **(j)** Expression of structure-guided SERBP1 mutants. Whole-cell lysates from cells expressing empty vector (EV), wild-type (WT) SERBP1 or the indicated mutants were analysed by immunoblotting with anti-SERBP1 and anti-FLAG antibodies. SERBP1 mutants targeting predicted GCN1 interfaces were F17A/D26A/K38A (GCN1-M), D337A/F359A (GCN1-N) and F17A/D26A/K38A/D337A/F359A (GCN1-N&M). The predicted GCN2 interface was targeted by S394A/D397A/D399A/D400A (GCN2-HRS), and the uS10-associated region by deletion of K346-G360 (DK346-G360). C-terminal FPxL motif mutants are indicated in the panel. **(k)** Structure-guided analysis of SERBP1 surfaces involved in anisomycin (ANS)-induced signalling. WT and SERBP1-KO cells complemented with EV, WT SERBP1 or the indicated mutants were treated with or without ANS. GCN2-dependent integrated stress response (ISR) activation was monitored by phosphorylated GCN2 and eIF2α, and ribotoxic stress response (RSR) activation by phosphorylated JNK and p38. SERBP1 and GAPDH were analysed as expression and loading controls. **(l)** Contribution of the SERBP1 C-terminal FPxL motif to ANS-induced signalling. WT and SERBP1-KO cells expressing EV, WT SERBP1 or the indicated FPxL mutants were treated with or without ANS. Phosphorylated and total GCN2 and eIF2α, JNK and p38, together with ZAKα, SERBP1 and GAPDH, were analysed by immunoblotting as indicated. **(m)** Association of GCN1 and GCN2 with disome fractions after SERBP1 complementation. Disome fractions from ANS-treated SERBP1-KO cells expressing EV, WT SERBP1 or the indicated GCN1-, GCN2- and uS10-interface mutants were analyzed by immunoblotting for GCN1 and GCN2. uS3 was used as a ribosomal marker, and SERBP1 expression was assessed to monitor complementation.

### SERBP1 determines the signalling output of collided ribosomes

Our mass spectrometry analysis indicated increased recruitment of RQC components to disomes upon SERBP1 loss (fig. S2c). We therefore next asked whether the GCN1-SERBP1 platform also regulates RQC engagement. ANS-induced uS3 ubiquitination was increased in SERBP1-KO cells (Fig. 4a), a phenotype reproduced by two independent SERBP1 siRNAs (fig. S7a). Ubiquitinated uS3 remained associated with ribosomal fractions (fig. S7b), indicating that SERBP1 loss enhances modification of ribosome-bound uS3. SERBP1 depletion also increased ubiquitinated forms of uS10 and eS10 in untreated cells and following ANS or azaC treatment (Fig. 4b, fig. S7d,e and table S6). These observations are consistent with our structural data showing that SERBP1 contacts uS10 and occupies the 40S head near eS10, potentially limiting access of ZNF598 to collided ribosomes.

To determine whether increased ribosomal ubiquitination was accompanied by enhanced RQC, we first analysed the V5-EGFP-K20-RFP-HA stalling reporter (*4, 32*) by immunoblotting (Fig. 4c). SERBP1 depletion increased the abundance of RQC intermediates (asterisks), which were abolished upon loss of ZNF598 or ASCC3. We then assessed the functional consequences of SERBP1 loss using the GFP-P2A-K20-P2A-RFP reporter, quantifying the RFP:GFP ratio by flow cytometry (Fig. 4d,e; fig. S7c). Thus, these findings confirm that SERBP1 restrains RQC activity. Structure-guided complementation further showed that the GCN1-M mutant restored the K20 reporter ratio to near-WT levels, whereas the GCN1-N and GCN1-N&M mutants, the uS10-region deletion and the FPxL mutants failed to restore the WT reporter ratio (Fig. 4d). These findings implicate distinct SERBP1 interaction surfaces in the regulation of the K20 reporter.

To determine whether this RQC-restricting function depends on GCN2 activation, we compared SERBP1-, GCN1- and GCN2-deficient cells. GCN1 deficiency phenocopied SERBP1 loss in the K20 reporter, whereas GCN2 deficiency had little effect (Fig. 4e). ANS-induced uS10 and eS10 ubiquitination was likewise enhanced in GCN1-KO cells but not to the same extent in GCN2-KO cells (Fig. 4f). These findings reveal a joint function for GCN1 and SERBP1 in restricting RQC that is separable from GCN2 activation. This distinction was also reflected in collision-associated complexes. Immunoblotting and mass spectrometry of disome fractions revealed genotype-dependent changes in GCN1, ZNF598 and other ISR-, RSR- and RQC-associated factors (Fig. 4g, fig. S7g and data S3). Broader proteomic analysis showed corresponding changes in translation- and RNA-surveillance factors without wholesale loss of ribosomal proteins (fig. S7f). Together, these data support a hierarchical model in which SERBP1 and GCN1 establish a signalling-competent collision state that restricts ZNF598-dependent RQC, while promoting GCN2 activation to trigger the ISR (Fig. 4h). Quantitative disome proteomics further showed that ZNF598 association was significantly increased in GCN1-KO cells following ANS treatment (1.949-fold relative to ANS-treated WT cells, P = 0.0211), whereas the difference under basal conditions was not significant (1.259-fold, P = 0.5929; fig. S7g).

**Fig. 4.**
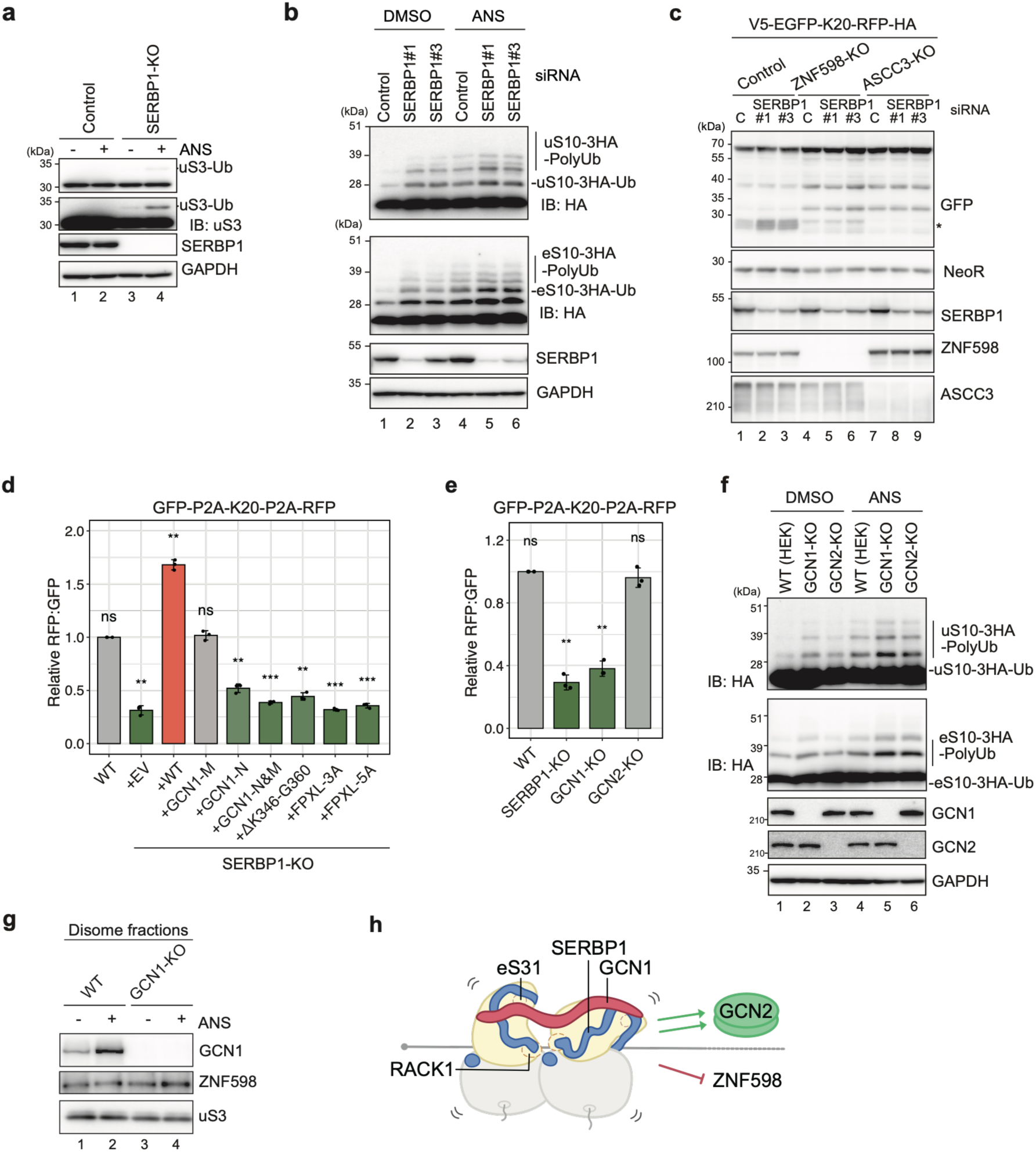
SERBP1 establishes the signalling identity of collided ribosomes. **(a)** Immunoblot analysis of uS3 ubiquitination in control and SERBP1-KO cells treated with or without anisomycin (ANS). Ubiquitinated uS3 (uS3-Ub) is indicated. SERBP1 and GAPDH were analysed as genotype and loading controls, respectively. **(b)** Ubiquitination of uS10 and eS10 following acute depletion of SERBP1. Cells expressing uS10-3HA or eS10-3HA were transfected with control siRNA or two independent SERBP1-targeting siRNAs (#1 and #3) and treated with DMSO or ANS. Ubiquitinated and polyubiquitinated forms of uS10-3HA and eS10-3HA were detected by immunoblotting with anti-HA antibodies. **(c)** Analysis of SERBP1-dependent processing of a defined ribosome-stalling substrate. Control, ZNF598-KO and ASCC3-KO cells expressing the V5-EGFP-K20-RFP-HA reporter were transfected with control or two independent SERBP1-targeting siRNAs (#1 and #3). Reporter products were analysed by immunoblotting for GFP. Asterisks indicate RQC intermediates. NeoR was used as an expression/loading control, and immunoblotting for SERBP1, ZNF598 and ASCC3 confirmed the indicated genetic perturbations. **(d)** Functional analysis of structure-guided SERBP1 mutants using the K20 stalling reporter. Relative RFP:GFP ratios are shown for WT cells and SERBP1-KO cells expressing empty vector (EV), WT SERBP1 or the indicated SERBP1 mutants targeting candidate GCN1- or ribosome-associated interaction surfaces. Values are normalized to WT. Bars show the mean ± SD; black dots represent individual samples (n = 3 per condition). Significance was assessed using a two-sided one-sample t-test against 1: **p < 0.01, ***p < 0.001; ns, not significant. **(e)** K20 reporter activity in WT, SERBP1-KO, GCN1-KO and GCN2-KO cells. Relative RFP:GFP ratios are shown normalized to WT. Bars show the mean ± SD; black dots represent individual samples (n = 3 per condition). Significance was assessed using a two-sided one-sample t-test against 1: **p < 0.01; ns, not significant. **(f)** Comparison of collision-induced ribosomal ubiquitination in WT, GCN1-KO and GCN2-KO cells. Cells expressing uS10-3HA or eS10-3HA were treated with DMSO or ANS, and ubiquitinated and polyubiquitinated species were detected by immunoblotting with anti-HA antibodies. **(g)** Association of collision-surveillance factors with disome fractions in WT and GCN1-KO cells treated with or without ANS. Purified disome fractions were analysed by immunoblotting for GCN1 and ZNF598, with uS3 serving as a ribosomal marker.| **(h)** Model for SERBP1–GCN1-dependent regulation of collision surveillance. SERBP1 forms a multivalent ribosome-associated platform with GCN1 that promotes GCN2 recruitment and activation, thereby triggering the integrated stress response. The SERBP1–GCN1 platform also restricts ZNF598-dependent ribosome quality control independently of GCN2 kinase activity.

## Discussion

Ribosome collisions activate multiple stress-signalling and quality-control pathways, yet how cells regulate these different outcomes remains unclear. Our findings distinguish the recognition of collided ribosomes from their acquisition of GCN2-signalling competence and identify SERBP1– GCN1 as a platform supporting the latter. This framework extends the concept that collisions function as a molecular gauge of translational stress (*10*) by suggesting that pathway selection depends not only on collision abundance or persistence, but also on the molecular machinery assembled on the collided ribosome. Rather than serving as a uniform signal, a collision can therefore acquire distinct functional states that favour stress signalling or quality control.

The identification of SERBP1 through azaC genetic screens places this function in the context of mRNA-damage-induced ribosome stress. AzaC incorporation into mRNA stalls elongating ribosomes and activates GCN1-GCN2-dependent ISR signalling, whereas RNF25-dependent eS31 ubiquitination restrains this response (*14*). Notably, the RNF25-targeted ubiquitination site on eS31 lies close to the SERBP1-binding region, suggesting a potential mechanism of allosteric regulation. The close genetic correlation of SERBP1 with GCN1 and GCN2, together with its requirement during ANS-induced elongation stress, argues that SERBP1 acts on a ribosomal state shared by mechanistically distinct perturbations. This interpretation aligns with an emerging view of GCN1 as a sensor of ribosomal state. Gcn1 spans stalled and colliding ribosomes (*11*), and reconstitution of the amino acid response showed that GCN1 acts as a di-ribosome sensor that recruits GCN2 and enables activating ribosomal interactions (*12*). Our findings add an additional organizational layer to this mechanism by showing that SERBP1 supports productive GCN1-GCN2 engagement, while restricting RQC jointly with GCN1 but independently of GCN2. Thus, the GCN1-SERBP1 platform functions not only as a GCN2 activator but also as an organizer of collision surveillance.

This view is reinforced by recent independent work showing that SERBP1 broadly regulates the mammalian ribosome interactome. Rosa-Mercado et al. found that SERBP1 loss increases association of ZNF598 and ASCC factors with polysomes without detectably increasing basal ribosome collisions, and demonstrated a protective role for SERBP1 against RNF10-RIOK3-dependent 40S degradation (*21*). These observations establish SERBP1 as a gatekeeper that limits inappropriate access of multiple surveillance pathways to the ribosome. Our results reveal a complementary positive function. SERBP1 promotes GCN2 recruitment and activation on collided ribosomes even when GCN1 association is retained. Thus, SERBP1 does more than limit surveillance-factor access; it helps establish a ribosomal state capable of productive ISR signalling. RACK1 may therefore act as a shared regulatory surface at which SERBP1 coordinates productive signalling with exclusion of competing surveillance machineries.

Several aspects of this model remain unresolved. It is not yet clear whether SERBP1 is constitutively positioned to receive GCN1 or is stabilized or rearranged following collision formation, and the candidate SERBP1-GCN2 interface requires direct structural validation. Different stresses may also generate collision states with distinct geometries or lifetimes, thereby shifting the balance among GCN1-GCN2, ZAKα, ZNF598 and RNF10/RIOK3 pathways. We therefore propose a hierarchical model in which collision formation, collision signalling and collision fate are separable steps. SERBP1-GCN1 converts a physical collision into a signalling-competent state, GCN2 executes the adaptive ISR, and failure to establish or maintain this state favours ubiquitin-dependent quality control (Fig. 4h). This framework reconciles the broad gatekeeping activity of SERBP1 with its specific role in GCN2 signalling and provides a molecular basis for how apparently similar ribosome collisions generate distinct cellular outcomes.

## Acknowledgments

We thank members of the Inada, Stingele and Beckmann laboratories for discussion and technical support. We thank Otto Berninghausen, Charlotte Ungewickell and Susanne Rieder for support with cryo-EM data acquisition. We thank Sylvia Mallok and Julia Philippou-Massier of the Gene Center genomics core facility (LAFUGA) for NGS services. We thank Steve Jackson for supporting the azaC CRISPR screens. We thank Jasmin Cartano and Amitkumar Fulzele (IMB Proteomics Core Facility) for excellent technical support. We thank the Human Genome Center at the University of Tokyo for providing computational resources. We thank Mario Dejung (IMB) for sharing computational expertise.

## Author contributions

Conceptualization, T.I., J.S. and R.B.; investigation, Z.Y., S.Y., T.D., T.S., T.T., C.M.E., M.T., S.L., S.H., C.K., A.L. and C.S.P.-C.; proteomic and genomic analyses, T.T., C.M.E., K.Y., Y.F., C.B., S.K., J.-X.C. and Y.S.; structural analysis, T.D., M.T. and R.B.; supervision, Y.S., R.B., J.S. and T.I.; writing - original draft, T.I. with input from J.S. and R.B.; writing - review and editing, all authors.

## Materials and Methods

### Cell culture and generation of knockout (KO) cell lines

HAP1 cells (C631; Horizon Discovery) were cultured in IMDM supplemented with 10% fetal bovine serum and 1% penicillin-streptomycin-glutamine. HEK293T cells (RCB2202; RIKEN BRC) were cultured in DMEM supplemented with 10% fetal bovine serum and penicillin-streptomycin (100 U ml-1) at 37 °C and 5% CO2. 293FT cells (R700-07; Thermo Fisher Scientific) were cultured in the same medium supplemented with 1× non-essential amino acids (11140-050; Thermo Fisher Scientific). SERBP1-, GCN1-, GCN2-, ZNF598-, RNF25-, ASCC3- and LTN1-deficient lines used in this study were generated by CRISPR-Cas9 genome editing or were derived from previously established lines, as indicated in the corresponding experiments. Clonal KO lines were validated by immunoblotting and, where applicable, by genotyping. Cells were routinely monitored for mycoplasma contamination. Exact culture media, supplements, clone identifiers and guide-RNA sequences are shown in table S8. FreeStyle 293-F suspension cells (Thermo Fisher) were grown in HyClone HyCell TransFx-H medium (Cytiva) supplemented with 100 U/mL Penicillin-Streptomycin (Gibco), 6 mM GlutaMAX (Gibco) and 0.01% Poloxamer 188 (Sigma Aldrich) at 37 °C, 5% CO2 and 80% humidity in a Multitron Cell (Infors HT).

### Plasmid construction

All recombinant DNA techniques were performed according to standard procedures using E. coli DH5α cells for cloning and plasmid propagation. Plasmids used in this study are listed in table S7. CSII-CMV-MCS-IRES2-Bsd (RDB04385), pCAG-HIVgp (RDB04394), and pCMV-VSV-G-RSV-Rev (RDB04393) were kindly provided by Dr. Hiroyuki Miyoshi (RIKEN BioResource Center, Ibaraki, Japan) (*33*). eS10-3HA or uS10-3HA lentiviral vectors were described previously (*34*). DNA cloning was performed by PCR amplification with gene-specific primers using PrimeSTAR HS DNA polymerase (#R010A, Takara-bio, Shiga, Japan) and T4 DNA ligase (#M0202S, NEB, Ipswich, MA 01938-2732, USA). Human SERBP1 cDNA was kindly provided by Dr. Yukihide Tomari. The SERBP1 mutants were generated using KOD mutagenesis Kits (TOYOBO, SMK-101). 3xFLAG tag sequences were included in PCR primers, and the amplified SERBP1 cDNA fragments including WT and mutants were inserted into multicloning site of CSII-CMV-MCS-IRES2-Bsd. pmGFP-P2A-K0-P2A-RFP and pmGFP-P2A-K(AAA)20-P2A-RFP were gifts from Dr. Ramanujan Hegde (Addgene plasmid #105686; http://n2t.net/addgene:105686; RRID: Addgene_105686) and (Addgene plasmid #105688; http://n2t.net/addgene:105688; RRID: Addgene_105688), respectively (*5*). Lentiviral vectors were constructed by subcloning GFP-P2A-K0/K20-P2A-RFP cassettes into multicloning site of CSII-CMV-MCS-IRES2-Bsd. To construct the pcDNA3.1(+) V5-EGFP-K20-RFP-HA vector, PCR-amplified V5-EGFP and RFP-HA fragments were sequentially inserted into the pcDNA3.1(+) vector (ThermoFisher V790-20). A DNA fragment encoding 20 consecutive lysine residues K(AAA)20 was then annealed and inserted between EGFP and RFP. For generating CRISPR KO cells, pSpCas9(BB)-2A-Puro (PX459) V2.0 (Addgene plasmid #62988; http://n2t.net/addgene:62988; RRID:Addgene 62988) (*35*) was linearized using BbsI (NEB). Guide RNA (gRNA) oligos were designed using CHOPCHOP (https://chopchop.cbu.uib.no/). Guide RNAs used in this study are listed in table S8. Reverse-complement oligonucleotides that encode guide RNA containing 5’ and 3’ overhangs for insertion were phosphorylated by T4 polynucleotide kinase (TAKARA), annealed and inserted into linearized PX459. All cloned DNAs amplified by PCR were verified by sequencing.

### CRISPR-mediated KO

For HEK293T KO generation, cells at approximately 50% confluence in 60-mm dishes were transfected with 2.5 µg PX459 guide-RNA plasmid using TransIT-LT1 (MirusBio, MIR2300). After 24 h, cells were replated at a 1:5 dilution and selected with puromycin (1 µg/ml) for 48 h. Surviving cells were isolated into 96-well plates, expanded for 10-14 days and screened by immunoblotting. KO clones were further assessed by genomic PCR and Sanger sequencing, using ISOGENOME (Nippon GENE, 314-08113) and KOD FX Neo (TOYOBO, KFX-201). The complete guide RNA sequences are provided in table S8.

### Lentivirus production and infection

293FT cells were grown to around 80% confluency on 6-cm plates. A transfection mixture containing 14.7 μL PEI-MAX reagent (CosmoBio, Koto, Tokyo, Japan), 1.2 μg pCAG-HIVgp (RDB04394), 1.2 μg pCMV-VSV-G-RSV-Rev (RDB04393) and 2.5 μg CSII-CMV-MCS-IRES2-Bsd (RDB04385) encoding WT or mutant SERBP1 or carrying the EGFP-P2A-K0/K20-P2A-RFP cassette was prepared in 500 μL Opti-MEM reduced-serum medium (Thermo Fisher Scientific, 31985-062). The mixture was incubated at room temperature for 25 min and gently added to the 293FT cells for 4 h incubation (37 °C, 5% CO2). The medium was then replaced with pre-warmed media (DMEM supplemented with 10% FBS, 1x NEAA). 48 hours after the start of the transfection, lentivirus-containing cell culture supernatants were collected and filtered (Millipore, MILLEX GV 0.45 μm). The resultant virus solution (4.5 mL) supplemented with polybrene (Sigma, H9268, 5 μg/mL) was added to HEK293T cells that were seeded at 4 × 10^5^ cells per 6-cm dish the day before infection. 24 hours after infection, cell culture medium was replaced with normal media. 48 hours after infection, cells were trypsinized and subjected to selection by blasticidin (10µg/mL) for 48 hours. Cells were then changed to normal media and passaged for experiments.

### Drug treatments

Cells were treated with anisomycin (MedChemExpress, HY-18982) or 5-azacytidine (Tokyo Chemical Industry, A2033) at the concentrations and for the durations indicated in the figures and legends. DMSO was used as vehicle control where appropriate.

### Genome-wide CRISPR-/Cas9 screens and correlation network analysis

Genome-wide CRISPR-/Cas9 screens were performed in WT HAP1, RNF25-KO and ZNF598-KO HAP1 cells using the Brunello lentiviral library (Addgene #73179), following the previously described screening protocol (*14*). Briefly, cells were transduced at a multiplicity of infection (MOI) of 0.25, maintaining a 500× coverage of the Brunello library throughout the screening workflow. After 24 h, cells were selected with 0.6 μg/mL puromycin for seven days, during which time they were divided into two replicates.

Seven days after selection, cells were treated with DMSO (control) or the respective IC50 concentration of azaC under chronic or acute treatment conditions. For chronic treatment, cells were passaged and re-treated every 48 h for a total of 10 days, with cell pellets collected on days 8 and 10 of treatment. For acute treatment, cells were exposed to a higher drug concentration for 48 hours and then passaged every 48 hours for a total of 10 days, with cell pellets collected on days 2, 6 and 10 of treatment. The concentrations used were as follows: WT cells, 240 nM for chronic treatment and 1.4 μM for acute treatment; RNF25-KO cells, 450 nM for chronic treatment and 0.8 μM for acute treatment; and ZNF598-KO cells, 650 nM for chronic treatment and 1 μM for acute treatment.

Genomic DNA was extracted from harvested cell pellets using the QIAamp Blood Maxi kit (Qiagen). The sgRNA cassettes were amplified by PCR using NEBNext Ultra II Q5 Mastermix (New England Biolabs) and i7 barcoded multiplexing primers (Merck) (table S10). PCR products were purified using a Qiagen PCR purification kit, quantified using the NEBNext Library Quant Kit (E7630), and multiplexed for sequencing on an Illumina NovaSeq 6000 system. Exorcise software was used to confirm CRISPR/Cas9 targets in GRCh38 (*36*). Guide RNA enrichment analysis was performed using DrugZ software to identify genes whose perturbation resulted in fitness effects between different conditions (*37*).

For correlation network analysis, normZ scores from the 15 genome-wide CRISPR screens across WT, RNF25-KO and ZNF598-KO HAP1 cells were analysed, including acute treatment time points (days 2, 6 and 10) and chronic treatment time points (days 8 and 10). Genes with identical or near-identical normZ scores across all 15 screens, reflecting sgRNAs that target more than one annotated gene, were collapsed into a single representative gene. The 250 genes with the highest absolute normZ score in any single screen were selected for pairwise Spearman correlation analysis. Pairwise Spearman correlation coefficients were calculated between gene score profiles across the screens, and gene pairs with an absolute correlation coefficient of |rho| > 0.7 were selected to construct the network. Genes with no correlation above this threshold were excluded, resulting in a network of 238 genes connected by 2,741 correlations. Correlation analysis was performed in R (v4.5.2) using igraph (v2.3.2).

### Cell-viability assays

HAP1 cells were seeded in 12-well plates and treated the following day with the indicated concentrations of azaC with or without 1 μM GCN2i (A-92; TargetMol, T11373), with treatment refreshed every 24 h. After 48 h, cell viability was measured using an AlamarBlue assay. Cells were incubated with 40 μg/mL resazurin (Sigma, R7017) in PBS for 1 h at 37 °C. Fluorescence was measured at 560 nm excitation and 590 nm emission using a Spark plate reader (Tecan). Data were normalized to the corresponding control within each replicate and are presented as mean ± SD of three biological replicates.

### Transfection of siRNAs or plasmids

Cells were transfected with siRNA using the reverse transfection method. Briefly, 50 pmol of siRNA was mixed with 4.5 μL of Lipofectamine RNAiMAX (ThermoFisher, 13778150) in 500 μL of Opti-MEM (Gibco) and incubated at room temperature for 15 minutes to allow complex formation. The siRNA mixture was then added to a 35 mm culture dish, followed by the addition of 2.5 mL of a cell suspension at a density of approximately 1 × 10⁵ cells/mL. For monitoring RQC activity, 4 ug of the pcDNA3.1(+) V5-EGFP-K20-RFP-HA vector was introduced into siRNA-introduced cells 24 hours after siRNA transfection using PEI-MAX reagent (CosmoBio, Koto, Tokyo, Japan). Cells were harvested for analysis 48–72 hours after siRNA transfection. Transfection conditions were scaled appropriately for different culture dish sizes and optimized based on cell type. siRNAs used in this study are listed in table S9.

### Immunoblotting

Primary and secondary antibodies are listed in table S11. For whole-cell lysates, cells were washed twice with PBS and lysed in RIPA buffer (20 mM Tris-HCl, pH 7.5, 150 mM NaCl, 2 mM EDTA, 10 mM NaF, 10 mM β-glycerophosphate, 1% Triton X-100, 0.1% SDS, 0.5% sodium deoxycholate and 40 mM N-ethylmaleimide). Lysates were cleared by centrifugation at 15,000 rpm for 10 min, mixed with sample buffer and heated at 95 °C for 5 min. Proteins (20 µg) were separated by SDS-PAGE, using MOPS-SDS buffer for HA-tagged uS10 and eS10 or Tris-glycine-SDS buffer for other proteins, and transferred to PVDF membranes (Millipore, IPVH00010). Membranes were blocked in 5% skim milk in TBS with 0.05% Tween 20 and immunoblotted using the antibodies listed in the accompanying materials table. Ubiquitinated uS3, uS10-3HA and eS10-3HA species were detected as indicated in the figures.

### OPP incorporation assay

HAP1 cells were seeded into 12-well plates and treated the following day with 5, 10 or 20 μM azaC for 4 h. Cells were then washed twice with PBS and incubated with 20 μM OPP for 30 min at 37°C. Cells were washed twice with PBS, trypsinized and transferred to U-bottom 96-well plates. After centrifugation at 300 x g for 3 min, cells were washed with 1% BSA in PBS and stained with eFluor780 viability dye (Thermo Fisher Scientific, 65-0865-14) for 20 min at 4°C in the dark. Cells were washed, fixed and permeabilized with CytoFix/CytoPerm (Thermo Fisher Scientific, 88-8824-00) for 20 min at RT in the dark. Cells were washed with Perm/Wash buffer and stored in 1% BSA in PBS at 4°C until further processing. Cells were spun down at 1700 rpm for 3 min and resuspended in freshly prepared click-chemistry mixture (39.5 mM Tris-HCl pH 8.0, 60 μM Alexa Fluor 488 azide, 4 mM CuSO4, 11 mM ascorbic acid and 10 nM DAPI) and incubated for 20 min at RT in the dark. Cells were washed twice with 1% BSA in PBS and finally resuspended in 1% BSA in PBS for flow cytometry. Samples were analyzed on a BD LSRFortessa (BD Biosciences) equipped with 405, 488 and 640 nm lasers, with at least 10,000 events collected per sample. Flow cytometry data were analyzed using FlowJo™ v10.7 software (BD Life Sciences). Fixable viability dye eFluor780 was used to exclude dead cells, and the mean fluorescence intensity (MFI) in the FITC channel was measured for live singlet cells. Values in the figures represent fluorescence intensities of live singlet cells normalized to the mean of the corresponding untreated control within each replicate. Data are presented as mean ± SD from three biological replicates.

### RNA-seq library preparation

HAP1 AAVS1, GCN2-KO, SERBP1-KO cells were seeded into 6-well plates and treated the following day with 20μM azaC for 4h. Cells were lysed in 500μL TRI Reagent (Sigma-Aldrich) and RNA was isolated using the Maxwell RSC 48 miRNA Tissue Kit (Promega) according to the manufacturer’s instructions. The quality of the isolated RNA was assessed using a NanoDrop ND-1000 spectrophotometer (Thermo Fisher Scientific) and a 2100 Bioanalyzer with the RNA 6000 Nano Kit (Agilent). High-quality RNA (RIN > 9) was used for library preparation. Libraries were generated using the CORALL mRNA-Seq V2 Kit (Lexogen) according to the manufacturer’s instructions and subjected to paired-end sequencing on an Illumina NextSeq system.

### RNA-seq data analysis

The samples were demultiplexed using an in-house tool based on their dual barcodes. Reads were aligned with STAR (version 2.7.11b) using the GRCh38 human reference genome and the associated gene annotations from Ensembl (GRCh38.p13). Reads per gene were counted using htseq-count (version 2.1.2) with settings --mode union and normalized using DESeq2 (geometric method, version 1.42.1). Further analysis was performed in R (version 4.3.2). Transcripts with at least 10 reads in at least three conditions were selected and Z-score normalized. High-confidence ATF4 target genes were selected and plotted as previously described (*22*).

### Phosphoproteome and ubiquitylome analysis after azaC treatment

Enzymatic protein digestion: HAP1 AAVS1 and SERBP1-KO cells were seeded in two 15 cm dishes per condition and treated or not with 5 µM azaC for 4 h. Cells were harvested by trypsinization and washed twice in PBS. Proteins were extracted in lysis buffer (3 M guanidinium chloride, 50 mM Tris-HCl pH 8.0, 1%[wt/vol.] CHAPS) at 90°C for 15 min. DNA was sheared for 3 min using a Covaris sonicator. Samples were processed using the SP3 approach (*38*). Briefly, 1 mg of proteins per sample were reduced in 5 mM DTT, alkylated in 15 mM chloroacetamide in the dark and quenched in 5 mM DTT. Enzymatic protein digestion was performed using trypsin (protein:protease ratio = 100:1) at 37°C overnight. Following acidification by formic acid, the peptides were purified by solid phase extraction in C18 cartridges.

Phosphopeptide enrichment: Phosphopeptide enrichment was performed in a microcolumn set-up using TiO2 (*39*). Desalted peptides were eluted directly in loading buffer (80% acetonitrile, 2% TFA, 1 M lactic acid (*40*)) and added to a microcolumn packed with 3 mg of TiO2 beads (Titansphere 5-µm; GL Sciences), followed by centrifugation at 100 × g. Afterwards, the microcolumn was washed once with loading buffer, followed by successive washes in buffer I (50% acetonitrile, 0.1% TFA) and buffer II (20% acetonitrile, 0.1% TFA) (*41*). Elution of the enriched peptides was performed twice by centrifugation at 500 × g in elution buffer (25% acetonitrile, 2% ammonia) directly into 45% formic acid. Thereafter, the acetonitrile was removed in a centrifugal evaporator at 45°C. The resultant peptide solution was further acidified by formic acid and purified by solid phase extraction in a C18 StageTip format (*42*).

Di-Gly-modified peptide enrichment: Desalted peptides were eluted in 50% acetonitrile. Afterwards, the acetonitrile was evaporated in a centrifugal evaporator. The resultant peptide solution was then adjusted with IAP buffer to reach 50 mM MOPS–NaOH (pH 7.2), 10 mM Na2HPO4, 50 mM NaCl (1X IAP buffer). Thereafter, the peptides were incubated with the PTMScan Ubiquitin Branch Motif (K-ε-GG) Immunoaffinity Beads (Cell Signaling) at 4°C overnight. Following sequential washes in 1X IAP buffer and water, peptides were eluted in 0.15% TFA and desalted in C18 StageTips.

Liquid chromatography tandem mass spectrometry: Peptides were separated via an in-house packed 45-cm analytical column (inner diameter: 75 μm; ReproSil-Pur 120 C18-AQ 1.9-μm silica particles, Dr. Maisch GmbH) on a Vanquish Neo UHPLC system (Thermo Fisher Scientific). Online reverse phase chromatography was performed through a 40-min gradient of 1.6-32% acetonitrile with 0.1% formic acid at a nanoflow rate of 300 nl/min. The eluted peptides were sprayed directly by electrospray ionization into an Orbitrap Astral mass spectrometer (Thermo Fisher Scientific). Mass spectrometry was conducted in data-independent acquisition (DIA) mode. Full MS spectra were acquired in the Orbitrap analyser in profile mode (scan range: 350 to 1,050 m/z; resolution: 120,000, target value: 3 × 10^6^, maximum injection time: 20 ms). The precursor ions were isolated for fragmentation at a window width of 4 Th over the range of 350 to 1,050 m/z. The fragment ions were recorded in centroid mode in the Astral analyser via higher energy collision dissociation (HCD; normalized collision energy: 26%, scan range: 150 to 2,000 m/z, target value: 5 × 10^4^, maximum injection time: 5 ms, default charge state: +2). Mass spectrometry data analysis: Raw data files were processed in DIA-NN software (version 2.2.0) (*43*) in library-free mode. An in silico predicted spectral library was generated in DIA-NN from the UniProt reference proteomes of H. sapiens (release 2024_03; 104,626 entries) and a list of common contaminants.

For the proteome data, the MS/MS spectra were searched against the predicted library using the following settings: Trypsin/P protease specificity, one missed cleavage allowed, maximum 1 variable modification, carbamidomethylation of cysteine as fixed modification, methionine oxidation as variable modification, peptide length range 7-30, precursor charge range 2-5, MBR and protein inference switched on, scoring in “peptidoforms” mode and “Genes” for proteotypicity, “RT-dependent” for cross-run normalization and “IDs, RT and IM Profiling” for library generation. The following settings were used to filter the identified precursors: Q.Value ≤ 0.01, PG.Q.Value ≤ 0.05, Lib.Q.Value ≤ 0.01, Lib.PG.Q.Value ≤ 0.01. Thereafter, the MaxLFQ algorithm (*44*) was applied to the “Precursor.Normalised” quantity using the DIA-NN R package for protein quantification. The LFQ intensities were further normalized by median-centering at the protein group level.

For the phosphoproteome data, the MS/MS spectra were searched against the predicted library using the following settings: Trypsin/P protease specificity, one missed cleavage allowed, maximum 3 variable modifications, carbamidomethylation of cysteine as fixed modification, methionine oxidation and phosphorylation at STY residues as variable modifications, peptide length range 7-30, precursor charge range 2-5, MBR and protein inference switched on, scoring in “peptidoforms” mode and “Genes” for proteotypicity, “Global” for cross-run normalization and “IDs, RT and IM Profiling” for library generation. The following settings were used to filter the identified precursors: Q.Value ≤ 0.01, PG.Q.Value ≤ 0.05, Lib.Q.Value ≤ 0.01, Lib.PG.Q.Value ≤ 0.01, Lib.Peptidoform.Q.Value ≤ 0.05. The detected phosphosites were then filtered to require a localization probability of at least 0.75. Thereafter, the “Precursor.Normalised” quantities were summed for each detected phosphosite. These summed intensities were normalized by median-centering across samples. Additionally, to correct for more missing values present in replicates 1 and 2 of the AAVS1 control group, the median intensity of each of the two replicates was adjusted separately by equalizing it to that of the intersect with replicate 4 as a reference.

For the ubiquitylome DIA data, the MS/MS spectra were searched against the predicted library using the following settings: Trypsin/P protease specificity, two missed cleavages allowed, maximum 3 variable modifications, carbamidomethylation of cysteine as fixed modification, methionine oxidation and di-Gly-K residues as variable modifications, peptide length range 7-30, precursor charge range 2-5, MBR and protein inference switched on, scoring in “peptidoforms” mode and “Genes” for proteotypicity, “Global” for cross-run normalization and “IDs, RT and IM Profiling” for library generation. The following settings were used to filter the identified precursors: Q.Value ≤ 0.01, PG.Q.Value ≤ 0.05, Lib.Q.Value ≤ 0.01, Lib.PG.Q.Value ≤ 0.01, Lib.Peptidoform.Q.Value ≤ 0.05. The detected di-Gly-K sites were then filtered to require a localization probability of at least 0.75. Thereafter, the “Precursor.Normalised” quantities were summed for each detected di-Gly-K site. These summed intensities were normalized by median-centering across samples and then used for downstream analysis.

Differential expression analysis was performed using the limma package in R (*45*). For each comparison, the proteins, phosphosites and di-Gly-K sites were filtered to retain only those detected in at least 3 out of the 4 replicates in either group of each comparison. Following imputation of the missing intensity values (except those missing in only 1 out of the 4 replicates), a linear model was fitted to assess the difference between the two groups, with adjustment for multiple testing using the Benjamini-Hochberg approach (*46*). The log2 fold change and the significance of the difference were displayed on a volcano plot. Only proteins, phosphosites and di-Gly-K sites with a minimum log2 fold change of 1 and an FDR-adjusted p value (q value) lower than 0.05 were considered differentially regulated.

Further computational analysis of phosphoproteomics data: To ensure changes reflected phosphorylation stoichiometry rather than protein abundance, phosphosite intensities were further normalized to their respective total protein levels. We employed PTM-SEA (v1.0.0) (*47*) utilizing the PTMSigDB reference dataset (v2.0.0) (*47*) to identify specific signalling features and estimate kinase activity based on kinase substrate enrichment analysis using integrated annotations from PhosphoSitePlus (PSP) (*48*) and the in-vitro Kinase-to-Phosphosite database (iKiP-DB) (*49*) which are integrated in the PTMSigDB reference dataset. Input identifiers were generated using flanking sequences (±7 amino acids) centered on the identified phosphosites to ensure uniqueness. PTM-SEA was executed on individual datasets using standard parameters with a minimum requirement of 5 kinase-substrate overlaps.

### Flow cytometry gating strategy for analysis of RFP:GFP expression

Representative plots illustrate sequential gating: FSC-A versus SSC-A to exclude debris, SSC-A versus SSC-H to select single cells, and FITC-A to identify GFP-positive cells. The RFP:GFP fluorescence-intensity ratio was determined within the GFP-positive population to quantify the polylysine-stalling reporter readout.

### Sucrose-density-gradient centrifugation and isolation of disome fractions

Cells were crosslinked with 0.5 mM dithiobis(succinimidyl propionate) for 15 min at room temperature and quenched with 25 mM Tris–HCl (pH 7.5). Cells were then washed with ice-cold PBS and lysed in SDG buffer (20 mM HEPES-NaOH [pH 7.5], 100 mM KCl, 10 mM MgCl₂, 1% Triton X-100, and protease inhibitor cocktail) and incubated on ice for 10 min. Cell lysates were centrifuged at 1,500 × g for 5 min at 4°C. Lysates containing 130 µg of total RNA normalized to 400 µL were treated with RNase A at 8 µg/mL for 10 min at room temperature. RNase digestion was stopped by addition of 200 U SUPERase·In RNase Inhibitor.

Sucrose buffer (10% or 40% sucrose in 20 mM HEPES-NaOH [pH 7.5], 100 mM KCl, and 5 mM MgCl₂) were prepared and stored at 4°C until use. 10%-40% Sucrose density gradients (SDG) were prepared using a Gradient Master 108 (BioComp). Cell lysates containing equal amounts of RNA described above were applied to the sucrose gradients and separated by centrifugation for 150 min at 230,000 ×g at 4°C by CP80NX (himac) ultracentrifuge and P40ST rotor (himac). The polysome profiles were generated by continuous absorbance measurement at 260 nm using a Piston Gradient Fractionator and TRIAX flow cell monitor FC-1-260 (BioComp) connected to a Micro Collector AC-5700 (ATTO) for fraction collection. Disome-region fractions were analysed by immunoblotting or DIA mass spectrometry.

### Immunoblotting of SDG fractions

Protein was precipitated from SDG fractions by addition of 10% volume TCA to each fraction. Samples were then incubated on ice for 20 min and centrifuged at 15,000 rpm at 4 °C for 15 min. Supernatants were removed and samples were washed once with 500 μL acetone, and centrifuged at 15,000 rpm at 4 °C for 10 min. After removing supernatants, samples were dried at room temperature for 10 min. Dried samples were resuspended in 2x sample buffer, then boiled at 95 °C for 5 min. Proteins were analyzed as described.

### DIA mass spectrometry of disome-associated proteins

For identification of disome-associated proteins shown in fig. S2b,c and fig. S7f,g, proteins recovered from isolated disome fractions were digested and analysed by data-independent acquisition (DIA) mass spectrometry. 100 ng of disome proteins were reduced with 10 mM DTT in 100 µL of 1% SDS at 70 °C for 10 minutes and alkylated with 15 mM iodoacetamide at room temperature for 15 minutes in the dark. Protein digestion was performed using a single-pot solid-phase-enhanced sample preparation (SP3)-based protocol on a KingFisher APEX system equipped with a 96 Combi head (Thermo Fisher Scientific) as previously described (*50*). Briefly, equal amounts of hydrophobic and hydrophilic Sera-Mag SpeedBead Carboxylate-Modified Magnetic Particles (# 65152105050250 and # 45152105050250, Cytiva) were washed with LC/MS-grade distilled water. Samples were mixed with an equal volume of 100% ethanol and conjugated to 40 µg of SP3 beads. Beads were then washed three times with 400 µL of 80% ethanol, and subjected to digestion in 0.5 µg of Trypsin Gold (#V5280, Promega) in 100 µL of 50 mM triethylammonium bicarbonate (# 90114, Thermo Fisher Scientific) for 3.5 hours at 37 °C. After digestion, beads were rinsed with 100 μl of LC/MS-grade distilled water. The eluate was combined with the digestion solution and acidified with trifluoroacetic acid at a final concentration of 0.1%, followed by desalting using Evotip pure columns (#EV2013, Evosep) according to the manufacturer’s instruction.

Peptides were analyzed by Evosep One (Evosep) coupled to an Orbitrap Fusion Lumos mass spectrometer (Thermo Fisher Scientific) equipped with a Dream Spray ING ion source (AMR). Analytes were separated on a C18 analytical column (Aurora ELITE [1.7 µm × 75-µm × 15-cm], IonOpticks) using the defined Whisper Zoom 20SPD gradient. Data were acquired using a data-independent acquisition (DIA) method with a FAIMS-pro set to −45 CV. Precursor MS spectra were collected in the range of 484–868 m/z (isolation window: 8 m/z, window overlap: 1 m/z). MS/MS spectra were collected in the range of 200–1,800 m/z at a resolution of 50,000. The HCD was set to 28. Raw files were analyzed using Proteome Discoverer 3.2 software (Thermo Fisher Scientific) with the Chimerys search engine against the Homo sapiens reference proteome (UniProt TaxID = 9606, version 2024-10-02). The mass tolerance for the fragment ions was set to 20 ppm. Methionine oxidation was set as variable modifications, and cysteine carbamidomethylation was set as a static modification for database searching. Maximum missed cleavage sites of trypsin were set to 2. Peptide identification was filtered at a false discovery rate (FDR) <0.01. Protein quantification was performed using the Fragment Ions Quantifier node in Proteome Discoverer 3.2. Protein-identification output tables for the DIA-MS analyses of disome-associated proteins are provided in data S2 (Fig. 2 and fig. S2) and data S3 (Fig. 4 and fig. S7).

### Ribosome-stalling reporter assays

Ribosome-associated quality control was assessed using the V5-EGFP-K20-RFP-HA reporter for immunoblotting and the GFP-P2A-K20-P2A-RFP reporter for flow cytometric analysis. Reporter constructs were expressed in control or KO backgrounds and, where indicated, combined with acute SERBP1 depletion. GFP-containing reporter products were analysed by immunoblotting, and NeoR served as an expression/loading control. For quantitative experiments, RFP:GFP ratios were normalized to WT.

### Flow Cytometry Analysis

For flow cytometric quantification of the stalling reporter, cells were washed three times in cold PBS and analysed on a CytoFLEX S (Beckman Coulter). Debris was excluded by FSC-A versus SSC-A gating, followed by singlet selection using SSC-A versus SSC-H. GFP-positive events were identified using FITC-A, and the RFP:GFP fluorescence ratio was calculated within the GFP-positive population. The gating strategy is shown in fig. S7c. At least 15,000 GFP-positive events were analysed per experiment using FlowJo. RFP:GFP ratios in different cell lines were normalized to WT, and the decrease of RFP:GFP ratio indicates enhanced ribosome stalling on the poly-lysine sequence while increased RFP:GFP ratio indicates readthrough.

### SERBP1 complementation and structure-guided mutagenesis

SERBP1-KO cells were complemented with WT SERBP1 or structure-guided SERBP1 mutants targeting predicted interaction interfaces. The GCN1-interface mutants were F17A/D26A/K38A (GCN1-M), D337A/F359A (GCN1-N) and F17A/D26A/K38A/D337A/F359A (GCN1-N&M); the GCN2-interface mutant was S394A/D397A/D399A/D400A (GCN2-HRS); and the uS10-region deletion removed residues K346-G360 (DK346-G360).

### In extracto cryo-EM sample and grid preparation

Cell extracts for cryo-EM samples were prepared essentially as previously described (*24*). FreeStyle 293-F cells were grown to 1.0 x 10^6^ cells/mL and treated with 100 µM ANS for 15 min. Then, cells were harvested by centrifugation at 250 x g for 5 min and washed once with 1x PBS containing 100 µM ANS. Cells were resuspended in Extract Buffer (20 mM HEPES/KOH pH 7.5, 150 mM KOAc, 5 mM MgCl2, 1 mM DTT, 1x protease-inhibitor cocktail (homemade), 0.05% octaethylene glycol monododecyl ether) and lysis was assisted by passing them through a 26-gauge needle five times. The crude lysate was cleared by centrifugation at 15,000 x g for 15 min. For grid preparation, 3.5 µL of cleared lysate were applied to glow discharged R3/3 300 mesh carbon support copper grids (QUANTIFOIL). The sample was incubated for 45 s on the grid before blotting for 3 s and plunge freezing in liquid ethane using a Vitrobot Mark IV (FEI) at 5 °C and 85% humidity.

### Cryo-EM data acquisition and processing

For the ANS treated in extracto sample, cryo-EM data was collected on a Titan Krios (FEI) equipped with a Falcon 4i director electron detector (Thermo Fisher) and a Selectris X energy filter (Thermo Fisher) at 300 kV and a pixel size of 0.727 Å/pixel. A defocus range of -0.5 to - 3.5 µm and a total electron dose of 40 e-/Å^2^ were applied. Movies were motion corrected using MotionCor2 (*51*) and initial contrast-transfer function (CTF) parameters were estimated using CTFFIND4 (*52*). In total, 43,174 micrographs were collected. For picking of ribosomal particles, crYOLO (*53*) (v1.7.6) was trained with manually picked particles from 200 randomly selected low-pass filtered micrographs. For training the PhosaurusNet architecture was used with default settings. The particles picked from the entire dataset (188,468 particles) were extracted in Relion (*54*) (v4.0.1) four times binned with a box size of 160 pixels. Extracted particles were transferred to CryoSPARC (*55*) (v4.6.0) and subjected to 2D classification. Good classes corresponding to 80S ribosomes were selected (133,659 particles) and subjected to an ab initio reconstruction followed by a homogeneous refinement. Then, a 3D classification without focus mask was performed to separate different translation states. Three classes corresponding to decoding 80S containing P/P and A/T tRNAs as well as eEF1A which made up the most abundant translation state (42%; 55,422 particles) were selected. These particles were imported back into Relion and re-extracted without binning at a box size of 640 pixels. The undecimated particles were again transferred to CryoSPARC for a final homogeneous refinement with global CTF and per-particle defocus refinement followed by local resolution filtering. This resulted in an overall resolution of 2.47 Å and revealed the presence of SERBP1 and ANS on these 80S particles. Then, local refinements for the 60S large subunit, 40S head and eEF1A with A/T tRNA were performed after using inverse masks for particle subtraction. This resulted in map local improvements with overall resolutions of 2.39 Å for the 60S subunit, 2.57 Å for the 40S head and 2.96 Å for eEF1A. For all three local refinements, local resolution filtering was performed. Finally, a composite map was generated using ChimeraX (*56*) (v1.12).

All cryo-EM maps were visualized using ChimeraX (*56*) (v1.12).

### Model Building

As initial model for our SERBP1-bound human ribosome the model PDB:8G6026 representative of the codon recognition step in decoding was used. The model was rigid body docked and then manually adjusted. For the interaction of RACK1 with the SERBP1 FPxL motif an AlphaFold 331 (AF3) prediction of full-length RACK1 and residues 387-408 of SERBP1 was used as starting model (fig. S6a and b). For the interaction SERBP1 with uS10 an AF3 prediction between full-length uS10 and SERBP1-324-408aa was used as a starting point (fig. S6a and b). For the SERBP1-40S beak interactions an AF3 multimer prediction using full-length eS31 and eS12 together with SERBP1-280-346aa was used (fig. S6a and b). For the interaction of the SERBP1 N-terminus with the 60S subunit another AF3 prediction with full-length eL30, eL27 and SERBP1-1-50aa was used (fig. S6a and b). All AF3 predictions were docked rigidly based on their ribosomal protein parts and manually adjusted. Sidechains were removed where local resolution did not allow unambiguous modelling. All manual adjustments were performed in Coot (*57*) (v9.8.96). Finally, the model was real-space refined using Phenix (*58*) (v2.2.1-6174) (table S1). All molecular models were visualized using ChimeraX (*56*) (v1.12).

### Quantification and statistical analysis

Biological replicate numbers and statistical tests are indicated in the corresponding figure legends where available. Quantitative data are presented as mean ± s.d. unless otherwise stated. Statistical significance was assessed using tests appropriate to the experimental design, with exact P values or significance thresholds reported in the final figures. No statistical method was used to predetermine sample size unless otherwise specified.

